# Exploiting Peptide Chirality and Transport to Dissect the Complex Mechanism of Action of Host Peptides on Bacteria

**DOI:** 10.1101/2025.09.24.678446

**Authors:** Siva Sankari, Markus F. Arnold, Vignesh M.P. Babu, Michael Deutsch, Graham C. Walker

## Abstract

Elucidation of the complex mechanisms of action of antimicrobial peptides (AMPs) is critical for improving their efficacy. A major challenge in AMP research is distinguishing AMP effects resulting from various protein interactions from those caused by membrane disruption. Moreover, since AMPs often act in multiple cellular compartments, it is challenging to pinpoint where their distinct activities occur. Nodule-specific cysteine-rich (NCR) peptides secreted by legumes, including NCR247, have evolved from AMPs to regulate differentiation of their nitrogen-fixing bacterial partner during symbiosis as well as to exert antimicrobial actions. At sub-lethal concentrations, NCR247 exhibits strikingly pleiotropic effects on *Sinorhizobium meliloti*. We used the L- and D-enantiomeric forms of NCR247 to distinguish between phenotypes resulting from stereospecific, protein-targeted interactions and those caused by achiral interactions such as membrane disruption. In addition, we utilized an *S. meliloti* strain lacking BacA, the transporter that imports NCR peptides into the cytoplasm. BacA plays critical symbiotic roles by reducing periplasmic peptide accumulation and fine-tuning symbiotic signaling. Use of the BacA-deficient strain made it possible to distinguish between phenotypes resulting from peptide interactions in the periplasm and those occurring in the cytoplasm. At high concentrations, both L- and D-NCR247 permeabilize bacterial membranes, consistent with nonspecific cationic AMP activity. In the cytoplasm, both NCR247 enantiomers sequester heme and trigger iron starvation in an achiral but BacA-dependent manner. However, only L-NCR247 activates bacterial two-component systems via stereospecific periplasmic interactions. By combining stereochemistry and genetics, this work disentangles the spatial and molecular complexity of NCR247 action. This approach provides critical mechanistic insights into how host peptides with pleiotropic functions modulate bacterial physiology.

**Author Summary:** Many organisms produce antimicrobial peptides (AMPs) to fight infections, but legumes have uniquely co-opted these molecules to control their symbiotic partners. During symbiosis between *Medicago truncatula* and *Sinorhizobium meliloti*, the plant secreted Nodule-specific Cysteine-Rich (NCR) peptides, transforms free-living bacteria into differentiated bacteroids that fix nitrogen but cannot reproduce outside the host. One such peptide, NCR247, exerts pleiotropic effects on the bacteria, acting on different subcellular locations, including membrane, heme, and proteins. Using a mirror-image (D-form) peptide, we disentangled peptide effects arising from generic physiochemical interactions versus stereospecific binding. The inner membrane protein BacA is known to play a protective role by importing NCR peptides into the cytoplasm. Using a bacterium lacking BacA, we were able to distinguish the effects of the peptide within and outside the cytoplasm. It was thought that BacA safeguards symbiotic bacteria by internalizing NCR peptides, thereby limiting their toxic membrane lytic effects, yet this has not been demonstrated. We show that BacA prevents lethal overstimulation of signaling pathways in the periplasm by internalizing the peptides. Our methods provide a framework for testing mechanism of action of new peptide-based antibiotics to combat multidrug-resistant bacteria.

## Introduction

Host secreted antimicrobial peptides (AMPs) constitute a crucial component of the mammalian innate immune system, characterized by their killing activity against pathogenic bacteria, fungi, viruses, and parasites. Beyond pathogen defense, host-secreted peptides also modulate symbiotic relationships [1,2,3]. One particularly interesting class of defensin-like peptides secreted by legume host plants, called the Nodule-specific cysteine-rich peptides (NCR), play an important role in modifying the life cycle of endosymbiotic bacteria [4]. The free-living, soil-dwelling bacteria enter developing nodules, chronically infect the plant cell, and endosymbiotically reside inside plant-derived symbiosomes. This symbiotic relationship is agriculturally, ecologically, and economically important. In the partnership between the Inverted Repeat Lacking Clade (IRLC) legume *Medicago truncatula* and *Sinorhizobium meliloti*, the free-living monoploid bacteria infect the plant, but the plant enslaves the bacteria, turning them into a polyploid, terminally differentiated endosymbionts, called bacteroids [4]. During this process, terminally differentiated bacteroids lose their ability to survive outside the host. The plant achieves this extreme control over the life cycle of bacteria by targeting bacteria with an arsenal of ∼700 small peptides known as NCR peptides, which contain conserved cysteine motifs. They drive profound physiological, transcriptional, and morphological changes underlying bacteroid differentiation, which include endoreduplication, altered membrane permeability, and cell elongation or branching. NCR production is tightly developmentally coordinated, with successive waves secreted into symbiosomes via an endoplasmic reticulum–dependent pathway requiring the DNF1 nodule-specific signal peptidase [5].

Approximately one-third of NCR peptides in *Medicago truncatula* are cationic [6] and, at high concentrations, exhibit potent antimicrobial activity against a range of bacteria and fungi [7,8]. In the nodule micro-environment, however, these peptides are non-lethal, instead modulating bacterial metabolism and altering their life cycle. NCR247, a well-studied NCR peptide, exerts complex effects on bacterial physiology through a combination of direct physicochemical interactions with bacterial cell envelopes and specific molecular engagements with intracellular targets [9]. These effects are often pleiotropic, simultaneously involving membrane disruption, modulation of signaling pathways [10], interference with translational machinery [11], and alteration of metal homeostasis [12]. In *Sinorhizobium meliloti*, BacA, an inner membrane protein critical for bacterial survival within host environments [13], facilitates the uptake of host-derived NCR peptides into the cytoplasm [14,15]. This is presumed to protect the bacterium from the peptides’ deleterious effects, thereby enabling intracellular persistence and symbiotic nitrogen fixation.

Many AMPs combat multidrug-resistant bacteria, making them promising candidates for therapeutic development [16]. While their classical activity involves pore formation or membrane permeabilization [17,18], increasing evidence shows that AMPs also translocate into cells and act on intracellular targets [19,20]. Intracellularly, they are shown to target vital biochemical processes, including DNA [21,22], RNA [23], and protein synthesis [24,25], enzymatic functions [26], and cell wall biosynthesis [27,28], highlighting their multifaceted mechanisms and potential to limit the development of resistance.

A significant obstacle to elucidating the precise mechanisms of NCR peptides and other antimicrobial peptides lies in resolving the multiplicity of subcellular compartments of the Gram-negative symbiotic bacteria—outer membrane, periplasm, cytoplasmic membrane, and cytoplasm—in which they exert their effects. Another difficulty lies in distinguishing whether their biological activities arise from non-specific interactions with membranes and small molecules or from specific protein–peptide engagements. This complexity is compounded by the capacity of these peptides to permeate bacterial barriers and modulate distinct physiological processes in various cellular locales, making it challenging to determine the contribution of each compartment and interaction type to the overall antimicrobial or symbiotic outcome. Disentangling chiral, target-specific interactions from generic achiral, charge-driven effects is crucial for the rational development of these peptides as antimicrobial agents.

Here, we address the complex mode of action of the nodule-specific cysteine-rich peptide NCR247 on the Gram-negative bacterium *S. meliloti* using two complementary experimental approaches. Stereospecificity analysis using D-NCR247 — We employed the D-enantiomeric form of NCR247, in which all amino acid residues are in the D-configuration, to distinguish phenotypes that arise from stereospecific protein–peptide interactions from those mediated by achiral membrane or small molecule binding effects. Subcellular action discrimination using a Δ*bacA* mutant — We utilized a *bacA* deletion mutant to differentiate between phenotypes resulting from NCR247 activity within the cytoplasm versus those occurring outside the cytoplasm, given the established role of BacA in peptide uptake and intracellular delivery. Through this dual approach, we aimed to parse chiral interaction–dependent effects from spatial determinants of peptide action, thereby refining our understanding of NCR247’s mechanisms under symbiotically relevant sublethal concentrations. While BacA is postulated to protect bacteria by sequestering peptides away from membrane targets through cytoplasmic internalization, the mechanistic basis for this protective role remains speculative, especially when NCR peptides are at sublethal concentrations inside the nodules. Our work here supports a specific role for BacA in shielding bacteroids under sublethal NCR247 exposure, a condition highly pertinent to the symbiotic state.

## Results

### Achiral Membrane Disruption by NCR247 at High Concentrations

Cationic antimicrobial peptides are known to induce visible alterations such as bleb and blister formation on the cell surface and, eventually, lysis [29]. L- and D-NCR247 are highly cationic peptides with a pI of 10.5. They also have a hydrophobic residue content of approximately 29%, which facilitates their insertion into microbial membranes. Their exceptionally high Boman index of 4.63 kcal/mol, which is the highest among NCR peptides, indicates a strong potential for binding to other proteins and membrane components due to hydrophobic interactions. This balance between hydrophobic and cationic residues is considered to underpin its membrane-disruptive function [30]. L-NCR247, when used at higher concentrations, has been shown to induce membrane blebbing and compromise membrane integrity in bacteria like *S. meliloti*, *Escherichia coli*, *Bacillus subtilis*, and *Salmonella enterica,* presumably due to non-chiral, direct interaction with the membrane [30,31]. To test if D-NCR247 exhibits the same phenotype on *S. meliloti*, we treated *S. meliloti* with 20 μM of L and D-NCR247 and performed a cell viability assay to determine toxicity. As expected, both L and D-NCR247 were equally bactericidal to the cells at higher concentrations (Fig 1A). Scanning electron microscopy (SEM) revealed comparable morphological features such as membrane blebbing, distortion, and cellular shrinkage in 15 μM peptide-treated samples (Fig 1B). This shows that at higher concentrations of the peptide, non-chiral interactions cause membrane destabilization, resulting in cell death. This aligns with the properties of typical antimicrobial peptides that rely on charge and amphipathy-based interactions with the bacterial membranes rather than specific recognition.

**Fig 1.**
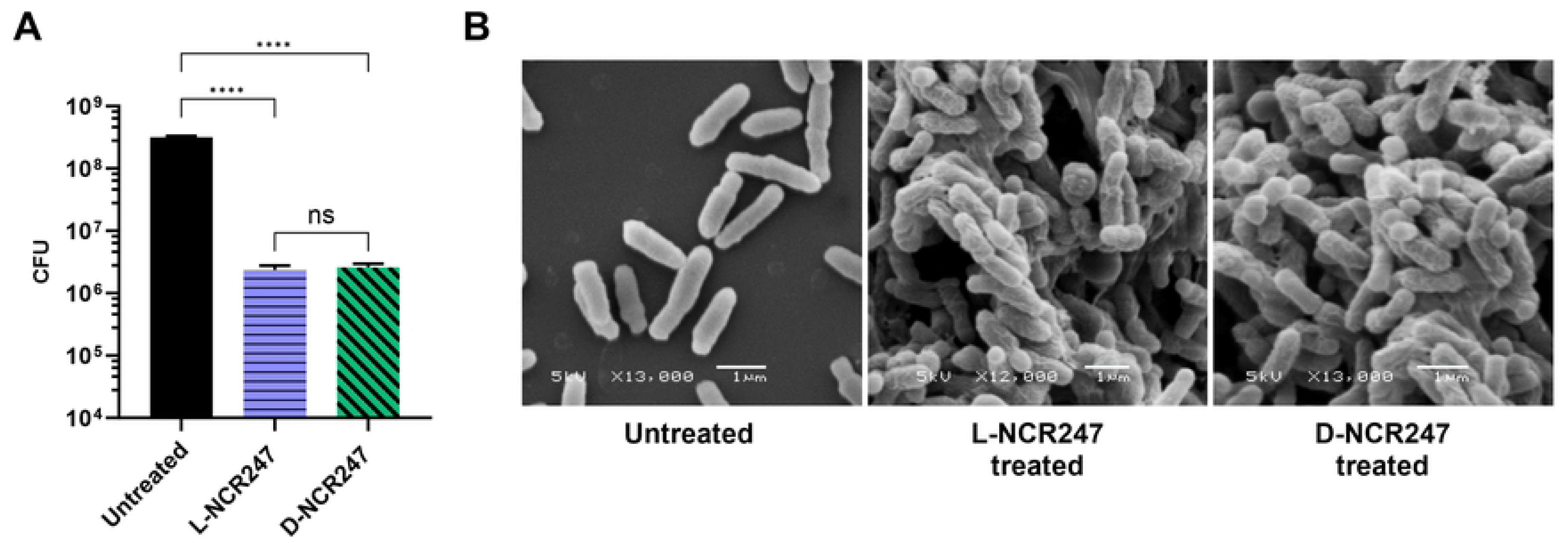
Non-chiral cell killing and membrane disruption effects. (**A)** A 100-fold reduction in colony-forming units when treated with 20 μM of L (blue) or D-NCR247 (green) when compared to untreated cells (black). (**B)** Scanning electron micrographs showing membrane blebs and blisters in 15 μM L- or D-NCR247 treated cells when compared to untreated cells. Data in A is presented as the mean of three biological replicates ± s.d. \*\*\*\**P* < 0.0001 versus untreated samples; two-way analysis of variance (ANOVA) with multiple comparisons. ns-no significant changes noted.

### L- and D-NCR247 binding to BacA

BacA has previously been shown to transport many varieties of peptides, including bleomycin [14]. The recent structure of SbmA, an *E. coli* homolog of BacA, reveals how the peptide binding cavity facing the periplasm in an open-outward conformation could allow the protein to promiscuously transport anti-microbial peptides [32]. Due to this promiscuity, it is reasonable to assume D-NCR247 will be transported through BacA, similar to L-NCR247. We tested whether D-NCR247 can bind purified SbmA with a similar binding curve to L-NCR247 in a Biolayer Interferometry experiment. Here, biotinylated L- or D-NCR247 was immobilized on a streptavidin sensor, and purified SbmA was tested for binding to the peptides. Both L and D-NCR247 bound to SbmA to the same extent (Fig 2A and 2B) further bolstering the idea that D-NCR247 could be transported by the same mechanism as L-NCR247.

**Fig 2.**
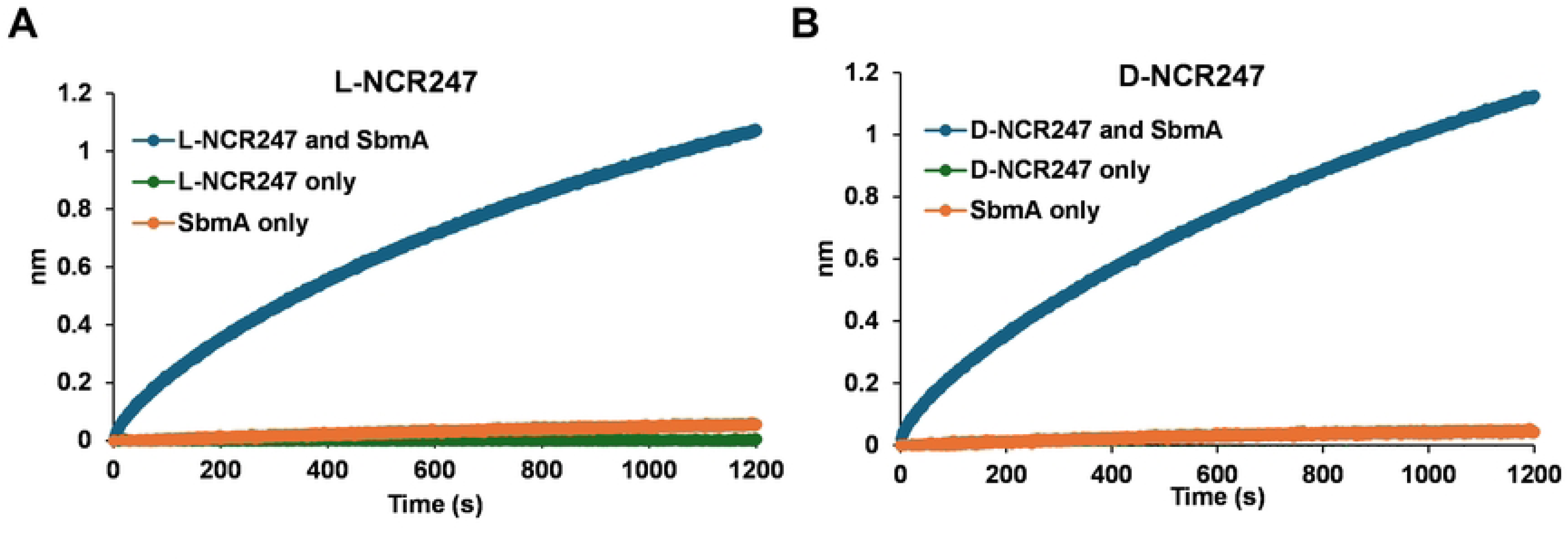
Binding of L and D NCR247 to SbmA (BacA homolog) Similar level of binding of L-NCR247 (**A**) and D-NCR247 (**B**) to SbmA in a biolayer interferometry experiment. Biotin-tagged L- or D-NCR247 was immobilized to the streptavidin sensor, and purified SbmA was loaded. Binding curve of SbmA interacting with the peptides is shown. L-, D-NCR247, and SbmA were loaded separately onto the sensors directly as controls to rule out non-specific binding to the sensor.

### Testing of symbiotically relevant phenotypes at sub-lethal levels of L-and D-NCR247

Next, we investigated the effects of sublethal concentrations of NCR247, which have previously been shown to modulate several physiologically pertinent processes to symbiosis [10]. To help us infer the underlying mechanisms, we employed both the L- and D-enantiomers of NCR247, as well as the Δ*bacA* mutant. This approach allowed us to systematically re-examine a variety of phenotypes previously attributed to NCR247, and to identify the contributions of chiral specificity and BacA-dependent uptake to these symbiotically relevant responses.

#### a) Chiral Activation of FeuP-FeuQ and ExoS-ChvI Two-Component Systems

Our previous microarray analyses examining the effects of sublethal concentrations of L-NCR247 on synchronized *S. meliloti* cultures revealed an extensive transcriptional response that included the upregulation of two regulons critical for symbiosis [10]. The FeuQ/FeuP regulon is controlled by a two-component response signaling system in which FeuP is the response regulator and FeuQ is an inner membrane protein with a short N-terminal cytoplasmic domain, a periplasmic sensor domain and C-terminal cytoplasmic histidine kinase response regulator domain [33]. Together they regulate genes, including *ndvA*, a cyclic beta-glucan exporter [33]. This regulon is usually activated under osmotic stress. The ExoS/ChvL regulon is involved in regulating multiple genes involved in exopolysaccharide production, motility, flagellar biosynthesis, and cell envelope integrity. ExoS has an overall structural organization similar to FeuQ with a periplasmic sensor and a cytoplasmic histidine kinase, while ChvL is the regulator [34,35,36]. s*mc01581* is a known direct target of ChvL [37]. To determine whether the activation of these regulons results from chiral interactions or from a general membrane stress response, we performed qRT-PCR analysis of representative genes from these regulons following treatment with 4 µM of either the L- or D-isoform of NCR247. As expected, genes from both regulons were increased in expression level upon L-NCR247 treatment. Transcript levels for multiple ExoS/ChvL-regulated genes, such as *smb21440* and *chvI* were significantly upregulated upon L-NCR247 treatment. In contrast, there was very little to no increase in gene expression in these regulons when the cells were treated with D-NCR247, showing that these signaling responses are due to specific chiral interactions of the peptide with proteins, rather than general membrane stress due to the cationic nature of the peptide (Fig 3A, 3B, and S1A and S1B Fig). To determine whether signaling due to chiral interaction occurs in the periplasm or inside the cytoplasm, we tested the same peptide treatments in a *ΔbacA* mutant. In the *ΔbacA* mutant, the increase in gene expression of these regulons was significantly higher than that of WT when treated with L-NCR247, indicating that there is increased accumulation of peptide in the periplasm, leading to overstimulation of the signaling pathways. In contrast, the D-NCR247 elicited minimal changes in gene expression and remained similar to untreated, confirming that the chiral interactions within the periplasmic space are responsible for the upregulation of FeuQ/FeuP and ExoS/ChvL two-component signaling by NCR247 (Fig 3A, 3B, and S1 A and S1B Fig).

**Fig 3.**
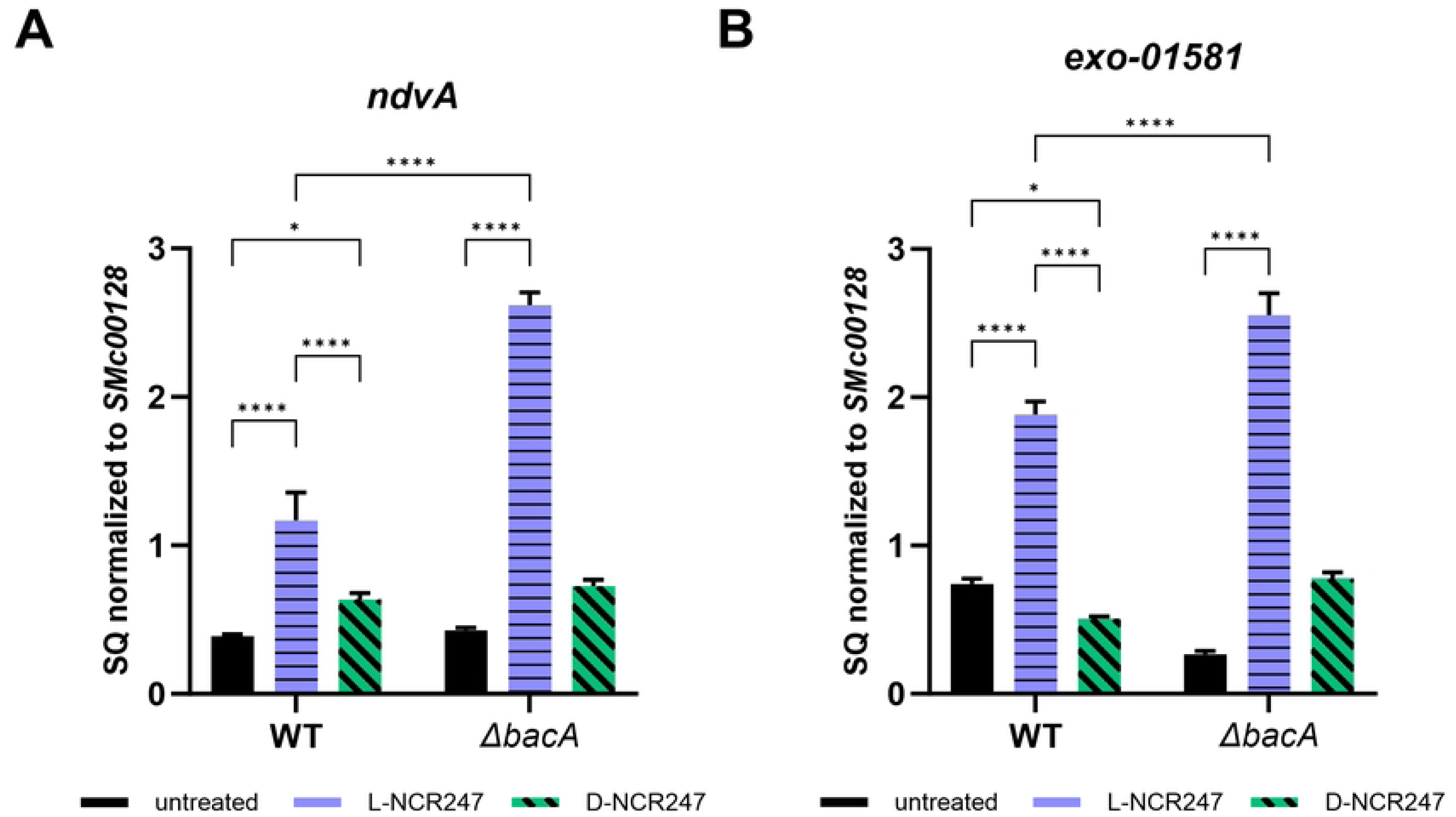
Chiral interactions in periplasm stimulate two-component signaling. Increase in expression of a gene target of FeuP/FeuQ signaling (*ndvA*) (**A**) and gene target of ExoS/ChvL signaling (*smc01581*) (**B**) upon treatment with 4 µM L-NCR247(blue) when compared to untreated cells (black) as quantitated by RT–qPCR analysis. Treatment with 4 µM D-NCR247 shows a significant reduction in expression in both cases (Green). In *ΔbacA* mutant there is significant increase in expression when treated with L-NCR247 when compared to wildtype treated at the same condition. The data are expressed as starting quantities (SQ) of respective mRNAs normalized to the control gene *smc00128* and are presented as an average of three technical replicates ± s.d, Two-way analysis of variance (ANOVA) with multiple comparisons was used to calculate P values.

#### b) Cell cycle regulation is only partially modified by chiral interactions

We then focused on the CtrA regulon, given its established role in controlling cell division and polarity [10,38]. As previously reported, treatment with a sublethal dose of NCR247 leads to expression changes in several genes of the CtrA regulon [10]. However, unlike the FeuP and ExoS regulons, the molecular components involved in signaling of this regulon are not well understood for CtrA. Gene expression analysis demonstrated that treatment with L-NCR247 led to a significant reduction in the expression of *ctrA* and gcrA in cell cycle regulation (Fig 4A and S2A). Upon D-NCR247 treatment, a more modest reduction in expression was observed that was nevertheless significantly different from the untreated samples. Thus, the responses associated with cell division inhibition are only partially dependent on the chiral interaction of the peptide.

**Fig 4.**
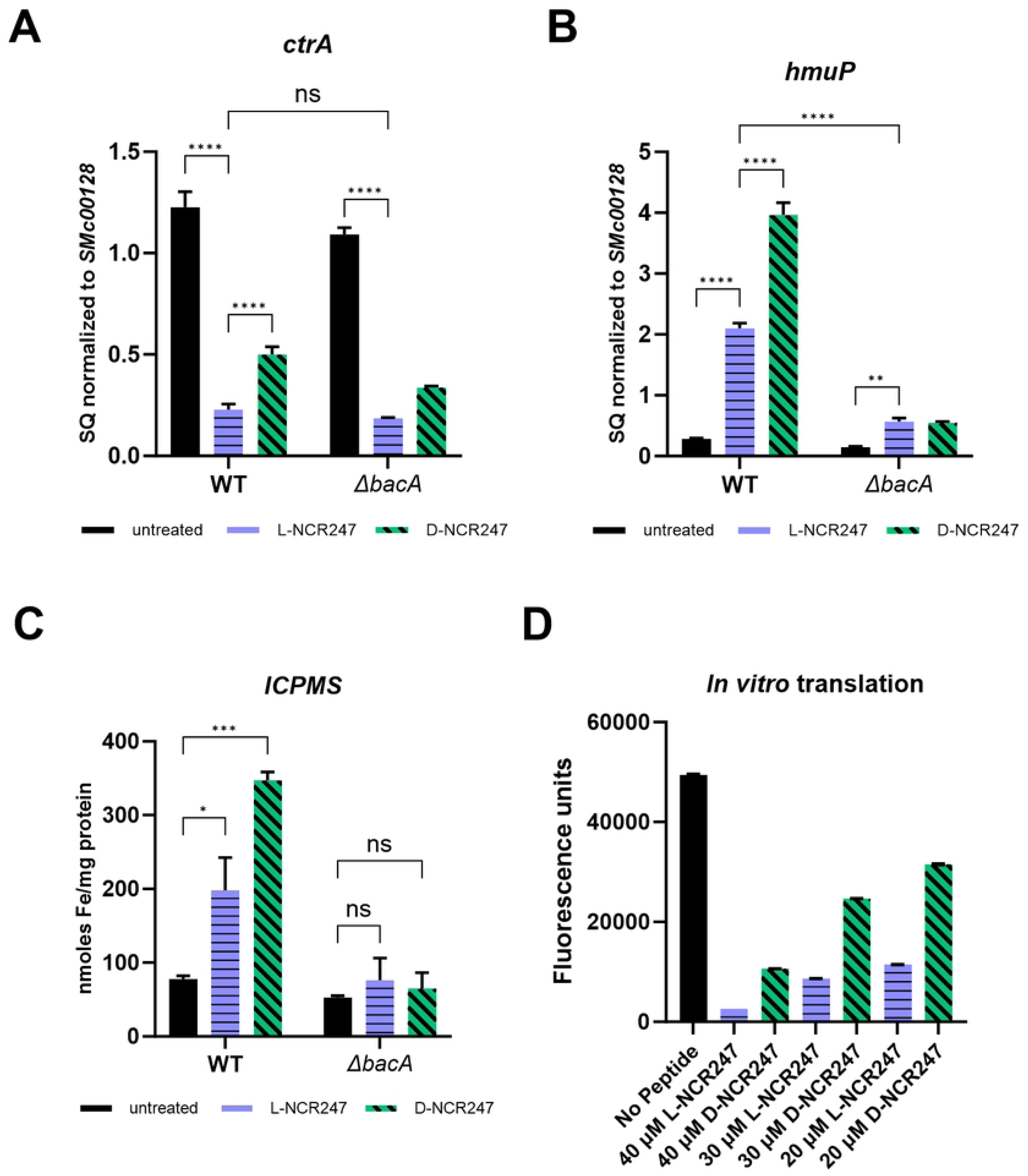
Phenotypes associated with peptide action in cytoplasm. **(A)** Decrease in expression of *ctrA* upon L-NCR247 (4 µM) treatment and partial recovery of expression upon D-NCR247 (4 µM) treatment as quantified by RT-qPCR analysis. Same treatments on *ΔbacA* mutant show a similar but modest response. (**B)** Increase in expression of heme import gene *hmup*, that is part of RirA regulon upon L-NCR247 treatment as analyzed by RT-qPCR analysis (blue) as compared to untreated samples(black). Further increase upon D-NCR247 treatment is noticed (green). Significant reduction of *hmup* expression upon both L-NCR247 and D-NCR247 treatment in *ΔbacA* mutant when compared to treatments on wildtype. For A and B, the data are expressed as starting quantities (SQ) of respective mRNAs normalized to the control gene *smc00128* and are presented as an average of three technical replicates ± s.d. (**C)** Increase in cellular iron content upon treatment with 4 µM of L-NCR247 (blue) and further increase upon treatment with 4 µM of D-NCR247 (green) in the wildtype, measured by ICPMS analysis. No significant change was observed in a *ΔbacA* mutant upon treatment when compared to the untreated. Data are presented as the mean of three biological replicates ± s.d. **(D)** Fluorescence measurement *(λ_ex_ = 485 nm, λ_em_ = 530 nm)* after treatment with lowering concentrations of L (blue) or D (green)-NCR247 in an *in vitro* translation assay using GFP vector. Data is presented as the mean of five independent reactions ± s.d. Two-way analysis of variance (ANOVA) with multiple comparisons was used to calculate P values.

To test whether this effect depended on cytoplasmic entry of the peptide, we repeated these experiments using a *ΔbacA* mutant strain. The responses due to treatment with L- and D-NCR247 were not as severe as compared to wildtype (Fig. 4A and S2A), indicating that the induction of the CtrA regulon and subsequent phenotypic effects are mediated only partially due to chiral interactions occurring in the periplasm. Since the responses are not apparent as in FeuP and ExoS signaling, there are likely some cytoplasmic mechanisms involved directly or indirectly in the control of the CtrA regulon.

#### c) Cytoplasmic Sequestration of Heme by NCR247 and induction of Iron Starvation Response is non-chiral

Next, we investigated the iron-related gene expression change controlled by the iron-responsive transcriptional regulator RirA. Reduced NCR247 binds heme with high affinity [12]. This sequestration of cytoplasmic heme deprives the cell of a critical iron source, thereby inducing genes involved in the iron starvation response. We have previously shown that the Δ*bacA* mutant is defective in inducing iron import genes, indicating that this heme sequestration happens in the cytoplasm [12]. We have biochemically shown that the D-NCR247 is capable of binding heme to the same extent as L-NCR247 [12]. Here, we tested whether D-NCR247 elicits the induction of genes involved in iron import similar to L-NCR247. In contrast to the two-component systems discussed above, when *S. meliloti* was treated with either L-NCR247 or D-NCR247, we observed upregulation of multiple iron acquisition and transport genes, including *hmup*, *entC*, and *fhup, which* was observed with D-NCR247 as well. As expected, this response was abolished in a *ΔbacA* mutant, showing that indeed iron-related responses are due to achiral interaction of NCR247 with heme in the cytoplasm (Fig 4B, S2B and S2C Fig). To confirm that the iron content increases upon an increase in expression of iron import genes, we measured cellular iron content using ICPMS following treatment with L- or D-NCR247. Similar to its effect on gene expression, D-NCR247 induces an increase in iron content of the cells when compared to untreated cells, and this response is greatly diminished in a *ΔbacA* mutant (Fig 4C). Interestingly, we notice that there is a significant increase in expression of iron uptake genes and iron content upon treatment with D-NCR247 compared to the increase observed with L-NCR247. This suggests that L-NCR247 may undergo either degradation or modification after entering the cytoplasm. To test this, we performed a pull-down assay with biotinylated L and D-NCR247 treated on cell cytoplasmic extracts. After incubation for 3 hours at room temperature, a silver stain of the pull-down clearly shows a band of the expected size (∼3 kDa) of NCR247 when D-NCR247 was used. Yet this band was not seen in extracts after the L-NCR247 pull-down. This suggests that there may be cellular factors, such as proteases, that act on L-NCR247 to modify or degrade the same (S2D Fig). This is important in the context of anti-microbial peptides, where BacA is suggested to play a role in protection against host peptides.

BacA is not only essential for peptide transport but also influences the modification of very long chain fatty acids (VLCFAs) in the inner membrane, affecting membrane stability. To distinguish these roles, a *ΔbacA* mutant complemented with the R389G BacA variant—defective in VLCFA modification but proficient in peptide uptake—showed restoration of iron-related phenotypes similar to wild-type BacA complementation. Structural studies of the homologous transporter SbmA by cryo-EM revealed an outward-open conformation with a conserved glutamate (E203 in SbmA, E207 in *S. meliloti* BacA) essential for proton binding and transport activity (36). Mutation of this residue (E207A) resulted in loss of BacA function and recapitulated the *ΔbacA* phenotype, underscoring the necessity of BacA’s peptide transport activity, rather than VLCFA modification, for mediating NCR247’s iron-related phenotypes (S2E Fig).

#### d) Inhibition of Translation by NCR247 Is Partially Chiral

Components of the protein translation machinery are the targets of some anti-microbial peptides[24,39]. Pull-down experiments with NCR247 have shown that many ribosomal proteins are capable of interacting with the peptide[9]. Additionally, previous *in vitro* translational assays have shown that L-NCR247 is capable of inhibiting translation[11]. To determine whether this inhibition is chirality-dependent, we performed *in vitro* coupled transcription-translation assays using GFP as a reporter. Reactions with L-NCR247 showed a dose-dependent reduction in GFP fluorescence intensity, indicating a reduction in protein synthesis. D-NCR247 treatment resulted in a reduction in translation efficiency, but the effect was smaller when compared to L-NCR247, suggesting that both chiral and achiral mechanisms may contribute to translational inhibition by NCR247(Fig 4D). We also confirmed these results by performing a western blot analysis against GFP which showed that changes in protein levels are comparable to the measurements using fluorescence (S2F and S2G Fig). These results support the model that, after NCR247 is internalized by BacA-mediated import, NCR247 may interact with cytoplasmic targets beyond heme, such as ribosomal components and RNA-binding proteins, in a manner that is partially chirality dependent. We had previously reported that one of the three regioisomers of disulfide cross-linked NCR247 was especially potent at inhibiting translation *in vitro*, and the cysteine residues are important for this activity [11].

### Mechanistic insights into protection of bacteroids by BacA using chiral peptides

BacA carries out two conceptually distinct critical roles in symbiosis. One role is to deliver certain NCR peptides into the cytoplasm where they can exert their critical intracellular symbiotic roles, for example, the induction of iron import via heme-sequestration in the case of NCR247. The other is to protect the endocytosed bacteria within the nodule cells from the potentially lethal anti-microbial peptide like effects of certain NCR peptides, including NCR247, by importing them into the cytoplasm, thereby reducing their concentration in the periplasm and the cytoplasmic membrane. A BacA deficient mutant must experience higher levels of NCR peptides in its periplasm than a wildtype during symbiosis, which can account for the extreme sensitivity of a Δ*bacA* mutant to cell death during symbiosis. This increased cell death of Δ*bacA* mutant cells has generally been attributed to increased membrane damage caused by the higher levels of NCR peptides in the periplasm, but this has not been tested to date.

A second possible death mechanism is suggested by: i) our observation of chirality-dependent periplasmic overstimulation of FeuP and ExoS signaling, and ii) a previous report that the absence of negative regulation of FeuP, which is mediated by FeuN, results in hyperactivation of FeuP/FeuQ signaling that leads to loss of *S. meliloti* viability. Connecting these two observations, we hypothesized that the hyperactivation of the two-component signaling caused by increased levels of NCR247 in the periplasm of the *ΔbacA* mutant may lead to detrimental effects on cell growth. To test this hypothesis, we compared the survival of wild-type and *ΔbacA* strains treated with sub-lethal doses of L- and D-NCR247 and observed growth over a period of 24 hrs. *ΔbacA* mutant exhibited inhibition of cell growth to L-NCR247 when compared to the wild type, even at a sublethal concentration of 4 μM. However, Δ*bacA* was less sensitive to treatment with the same concentrations of D-NCR247 when compared to the wildtype (Fig 5A and 5B). This suggests that in *ΔbacA*, when NCR247 cannot enter the cytoplasm, the peptide accumulating in the periplasm is detrimental to the cell. These findings may explain the activity of NCR247 even at very low, sublethal concentrations. Additionally, this inhibition of cell growth is due to the stereospecific interaction of the peptide with a periplasmic or outer membrane protein, since this inhibition is lost during D-NCR247 treatment. We then tested the cell viability at an intermediate concentration of 8 μM. While the cell viability of wildtype was reduced to the same extent with treatment of L- and D-NCR247, in a *ΔbacA* mutant, treatment with L-NCR247 is significantly more bactericidal than treatment with D-NCR247 (Fig 5C). This suggests that overaccumulation of L-NCR247 outside the cytoplasm imposes a severe bactericidal activity in a *ΔbacA* mutant in addition to the non-chiral toxicity due to membrane alterations. In contrast, the cell growth inhibition caused by D-NCR247 on wild-type cells can be recovered by the addition of increasing concentrations of iron (Fig 5D,5E and 5F). This suggests that both L- and D-NCR247 bind to heme in the cytoplasm and cause a bacteriostatic effect that can be recovered at high concentrations of iron.

**Fig 5.**
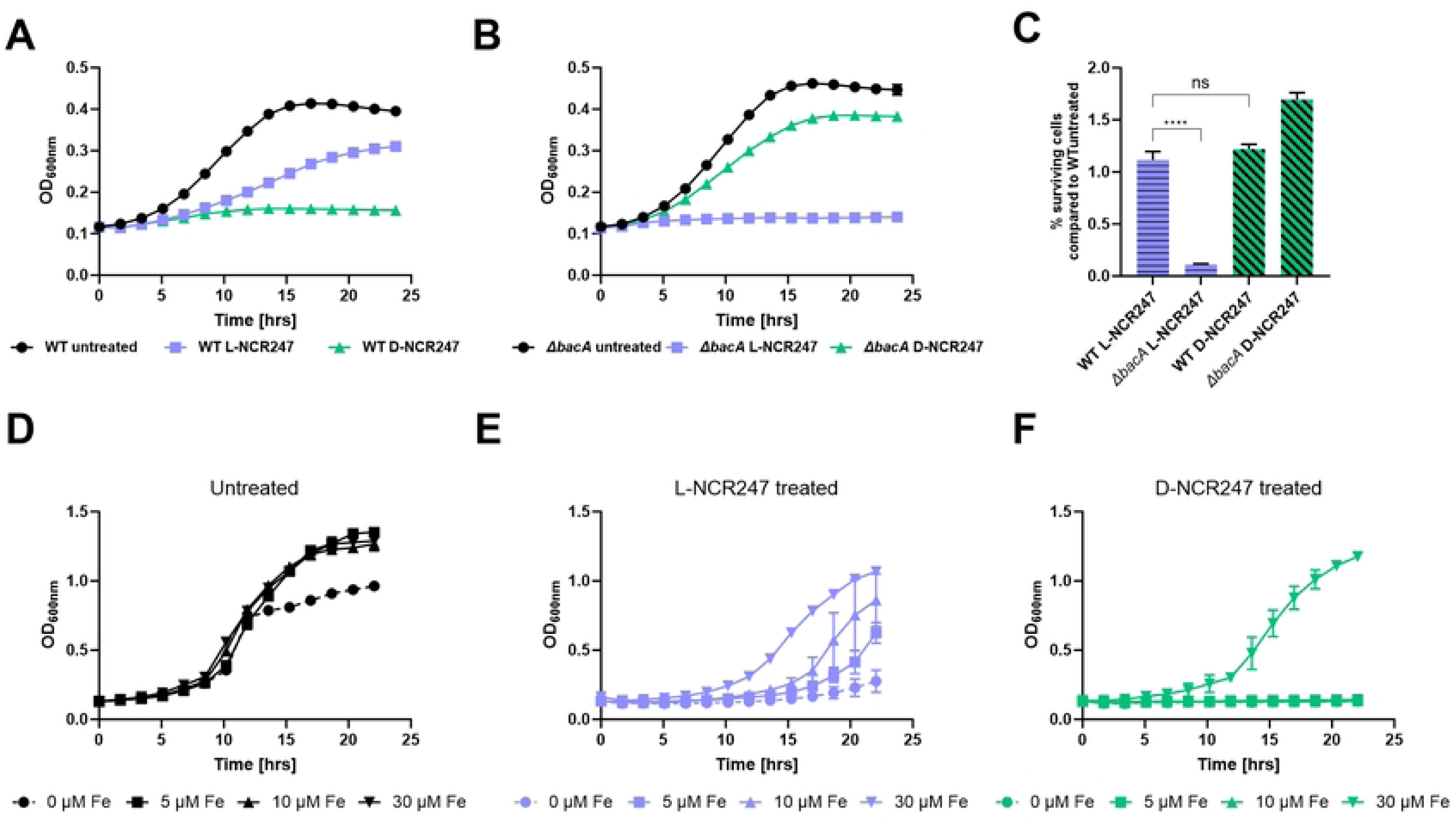
Accumulation of peptide in periplasm leads to reduction in cell growth of *ΔbacA* mutant. **(A)** Growth curve analysis over a period of 24 hours on wildtype cells treated with 4 µM L- or D-NCR247 showing increased growth inhibition upon treatment with D-NCR247. **(B)** Growth curve analysis over a period of 24 hours on *ΔbacA* mutant cells treated with 4 µM L- or D-NCR247 showing increased growth inhibition upon treatment with L-NCR247. **(C)** Cell viability assay of 8 µM L- and D-NCR247 treated wildtype and *ΔbacA* mutants. After treatment, cells were serial diluted, spotted and colonies were counted. % survival cells when compared to wildtype untreated cells are plotted. **(D, E and F)** Growth curve analysis of wildtype cells grown in 0 μM (●), 5 μM (▪), 10 μM (▲) or 30(▼) μM FeSO_4_ (D), upon treatment with L-NCR247 (E) and D-NCR247 (F). Data is presented as the mean of three biological replicates ± s.d.

We then investigated the mechanism of how the NCR247 peptide’s chiral interactions in the periplasm might mediate the cell death caused by periplasmic overaccumulation of NCR247 in a *ΔbacA* mutant. In the FeuP/FeuQ two-component relay system, FeuN is a periplasmic protein that prevents hyperactive signaling by directly binding and negatively regulating FeuQ[40]. Since our results indicated that hyperactivation of FeuP/FeuQ signaling happens during L-NCR247 treatment on *ΔbacA*, we hypothesized that abolishing this signaling response might restore the cell viability of this mutant. To test this, a *ΔbacA ΔfeuP* double mutant was created. Consistent with our hypothesis, in a *ΔbacA ΔfeuP* double mutant, the lethality due to the absence of BacA when treated with L-NCR247 is greatly diminished. This is noticed both at a sublethal concentration in a growth curve analysis (Fig 6A and 6B) and at a moderate concentration in a cell viability assay (Fig 6C). This shows that BacA plays a vital role in fine-tuning the amount of peptide available for stimulating signaling in the periplasm at physiologically relevant concentrations. This is also the direct clue for defining the role of BacA during symbiosis, where peptide concentrations are not lethal, yet enough to trigger signaling cascades.

**Fig 6.**
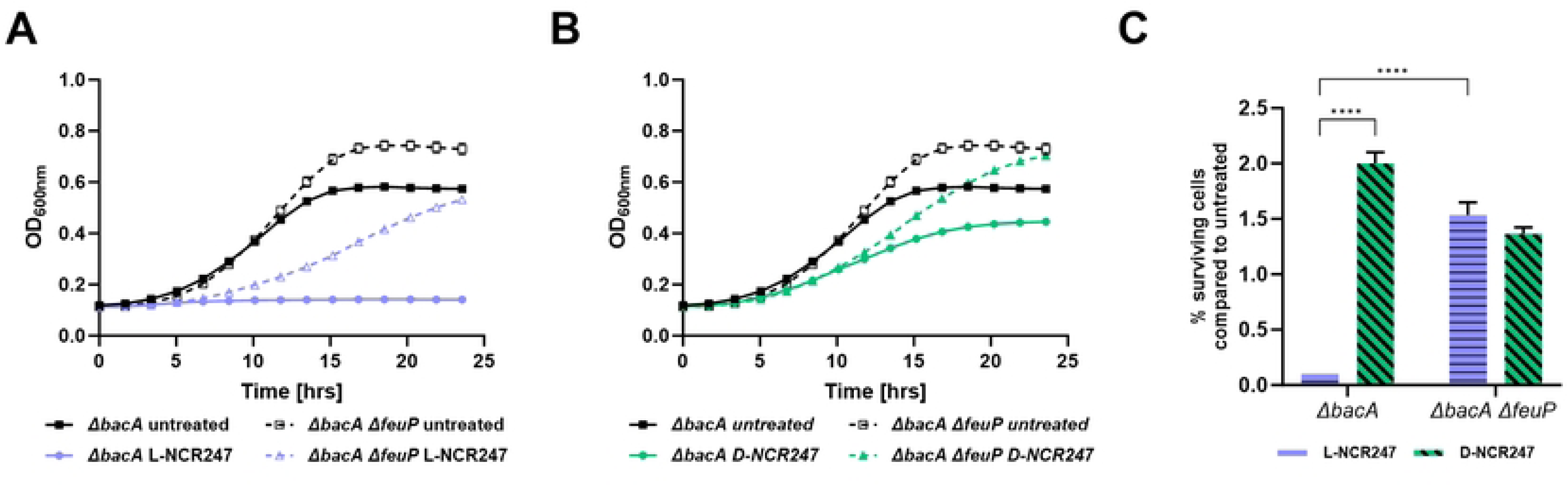
Deletion of two component signaling through FeuP in a *ΔbacA* mutant reduces growth inhibition by L-NCR247. Growth curve analysis of *ΔbacA* and *ΔbacA ΔfeuP* treated with 4 µM L-(blue) **(A)** or D -(green) (**B**) NCR247. **C**. Cell viability assay of 8 µM L- and D-NCR247 treated *ΔbacA* and *ΔbacA Δfeup* mutants. After treatment, cells were serial diluted, spotted and colonies were counted. % survival cells when compared to wildtype untreated cells are plotted. Data is presented as the mean of three biological replicates ± s.d.

To test whether the cysteines play a role in the toxicity associated with chiral interactions of L-NCR247 outside the cytoplasm, we used L and D-NSR247, in which serines replace the cysteines. NSR247 is shown to have no toxicity up to 20 μM. However, at 20 μM, while the WT is sensitive to both L and D-NSR247, the *ΔbacA* is fully resistant to both the peptides (S3A and S3B Fig). This shows that overaccumulation of NSR247 does not cause the lethal phenotype as NCR247, indicating that disulfide bonds are necessary for the lethal phenotype associated with NCR247 accumulation outside the cytoplasm. We have previously shown that NSR247 is capable of non-specifically binding to heme with very low affinity. The susceptibility of wildtype to L and D-NSR247 at these high concentrations may be due to weak binding of NSR247 to heme or other proteins in the cytoplasm. To confirm the phenotypes associated with *ΔbacA mutant* are not directly related to the VLCFA defects, we performed the growth curve analysis using L- and D-NCR247 on ΔbacA mutant complemented with wildtype copy of *bacA* or the R389G mutant version of *bacA*. No difference was noted between the two strains confirming the phenotypes are not due to VLCFA modification (S3C Fig).

### Conservation Across Diverse Host-Associated Bacteria

Homologues of BacA are present in many pathogenic bacteria. BacA is shown to be essential for chronic infection in mice infected with *Mycobacterium tuberculosis* [41] and *Brucella abortus* [42], indicating its importance for long-term bacterial survival inside the host. BacA from *M. tuberculosis* is shown to partially complement the lethality of the *S. meliloti ΔbacA* mutant upon peptide treatment. And a *ΔbacA* mutant in *B. abortus* shows modifications in VLCFA [13]. We investigated whether BacA from M. tuberculosis and B. abortus exhibits similar phenotypes as S. meliloti BacA towards L- and D-NCR247. BacA from *M. tuberculosis* and *B. abortus* follow a similar pattern in response to both the peptides, albeit to different extents (Fig 7A,7B and 7C). This suggests that the findings can be translated to the context of host-pathogen interactions and how these pathogens tolerate anti-microbial peptides from their respective hosts. This suggests a conserved, evolutionary role in enabling bacteria to survive hostile, antimicrobial peptide-rich environments.

**Fig 7.**
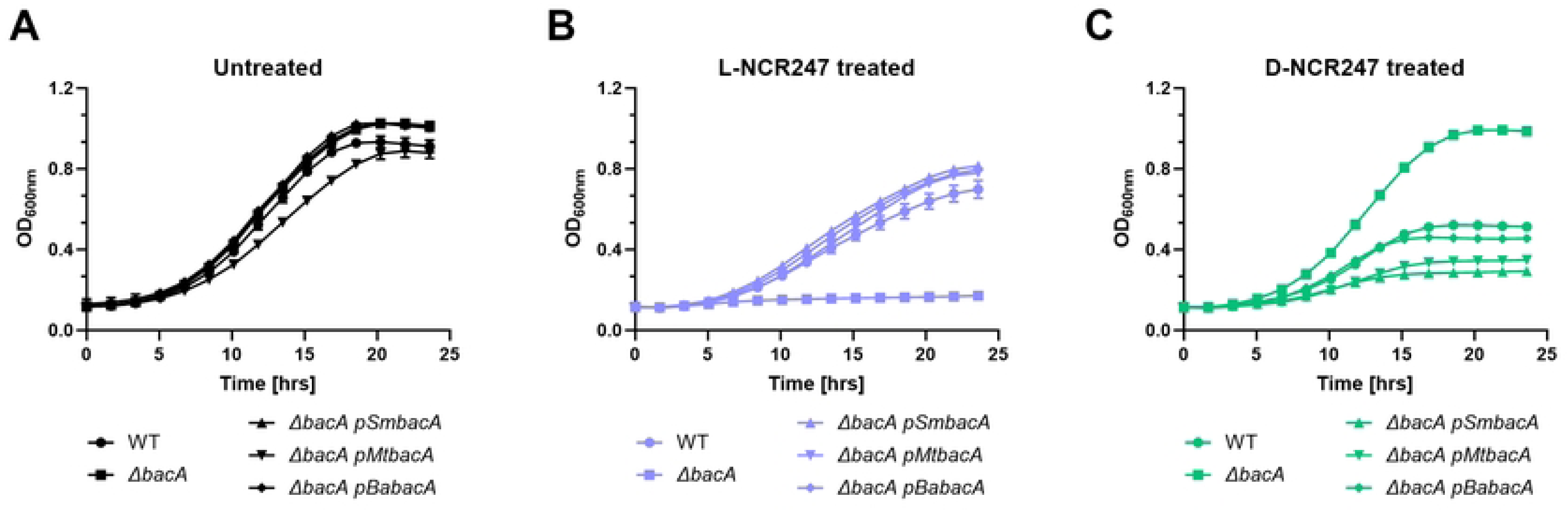
Complementation of BacA phenotype with homologs from pathogens. Growth curve analysis was performed on wildtype or *ΔbacA* mutant complemented with *Sinorhizhobium meliloti* BacA (pBacA), or BacA homologs of *Mycobacterium tuberculosis* (*pMtBacA*) or *Brucella abortus (pBaBacA)*. Untreated cells are shown in **(A)**, L-NCR247 treatment in **(B)** and D-NCR247 treatment in **(C).** Data is presented as the mean of three biological replicates ± s.d.

## Discussion

Our study provides new mechanistic insights into how the nodule-specific cysteine-rich peptide NCR247 exerts its effects on *S. meliloti*. By systematically combining stereoisomeric forms of NCR247 with a *ΔbacA* background, we were able to discriminate between (i) achiral, charge-driven membrane disruption, (ii) stereospecific, protein-dependent interactions within the periplasm and cytoplasm, and (iii) BacA-dependent modulation of subcellular exposure. This integrative approach has clarified several unresolved aspects of NCR peptide biology and of BacA functionality during symbiosis. S4 Table shows the summary of the findings.

At high concentrations, both L- and D-enantiomers of NCR247 destabilized bacterial membranes, leading to similar bactericidal outcomes. This is consistent with the behavior of conventional cationic AMPs, whose activities are primarily dictated by physicochemical parameters, such as net positive charge, and interaction with negatively charged bacterial envelopes. Importantly, this achiral mode of action explains the antimicrobial capacity of many NCRs when tested *in vitro* at elevated peptide doses. However, such concentrations are unlikely to be physiologically relevant in the nodule environment, suggesting that peptide-mediated symbiotic regulation involves more subtle, molecularly specific mechanisms.

At sublethal concentrations, only L-NCR247 elicited robust induction of FeuQ/FeuP and ExoS/ChvI two-component systems, while the D-NCR247 did not. This demonstrates that NCR247 engages in chiral interactions with periplasmic sensory proteins, triggering signaling cascades that alter envelope physiology, exopolysaccharide production, and stress responses. The stereospecific activation of these systems underscores a protein-targeted mechanism that could allow the host plant to fine-tune bacteroid physiology through evolutionarily selected peptide structures. In wild-type *S. meliloti*, BacA-mediated import redirected NCR247 into the cytoplasm, both mitigating excessive periplasmic signaling and enabling cytoplasmic phenotypes such as heme sequestration and translational inhibition. In the *ΔbacA* mutant, however, L-NCR247 accumulated in the periplasm, resulting in toxic hyperactivation of signaling pathways and growth arrest, even at nonlethal concentrations. Intriguingly, D-NCR247, which fails to engage in stereospecific protein interactions, accumulated without such detrimental outcomes, emphasizing that toxicity under these conditions is explicitly linked to chiral periplasmic binding events rather than nonspecific peptide accumulation. Reduction of toxicity due to L-NCR247 treatment in a *ΔbacA ΔfeuP* double mutant strongly bolsters this idea. Our findings therefore provide direct evidence to the earlier hypothesis that suggested BacA provides a generic protective function by internalizing toxic peptides. Our work further refined the model, showing that BacA appears to play a more sophisticated role in buffering the periplasmic environment against hyperactivation of regulatory systems, while simultaneously allowing controlled cytoplasmic engagement of peptides with intracellular targets. This advances a mechanistic framework for how BacA enables long-term symbiotic persistence despite continuous exposure to NCR peptides.

Inside the cytoplasm, both enantiomers were able to sequester heme, leading to an iron starvation response. The equivalence of L- and D-NCR247 in this activity confirms that heme binding is an achiral interaction based on physicochemical complementarity. This strongly supports prior biochemical data showing peptide–heme binding. The reliance on BacA for this phenotype further underscores its essential role in peptide transport across the inner membrane. Translational inhibition, by contrast, represented a mixed phenotype. While both enantiomers reduced protein synthesis, the more pronounced effects of L-NCR247 suggest contributions from stereospecific binding to ribosomal or translational factors. Thus, peptide functionality in the cytoplasm is stratified into achiral (heme sequestration) and partially chiral (translation inhibition) activities, once again highlighting the mechanistic complexity of NCR action.

Our experiments with NSR247 variants, in which serines replaced cysteines, pointed to an essential role for disulfide bonding in chiral, periplasmic signaling and associated toxicity. The absence of lethality in *ΔbacA* backgrounds with NSR247 treatment suggests that the structurally constrained, disulfide-stabilized form of NCR247 is required for effective engagement with periplasmic protein partners. This provides molecular insight into why evolution has favored cysteine-rich motifs across the NCR family and related defensin-like peptides, enabling them to adopt precise folds necessary for target recognition. Homologs of BacA in pathogens such as *M. tuberculosis* and *B. abortus* complement aspects of the *S. meliloti ΔbacA* phenotype, highlighting a conserved peptide-handling strategy that supports bacterial persistence in diverse hosts.

Our findings highlight how plants exploit the chemical versatility of AMPs not only for defense, but also to remodel microbial physiology in prolonged mutualistic associations. Future work should aim to identify the precise periplasmic binding partners responsible for NCR247’s stereospecific effects. These results provide critical insights for the rational design of peptide-based antimicrobials. Identifying the precise balance between achiral, charge-driven membrane disruption and stereospecific, protein-targeted effects could enable the design of peptides that minimize off-target immune responses while enhancing specificity for microbial proteins.

## Materials and Methods

### Growth conditions

All strains of *S. meliloti* were routinely grown in LB medium supplemented with 2.5 mM CaCl_2_ and 2.5 mM MgSO_4_ (LBMC) in the presence of 200 μg ml–1 streptomycin at 30 °C for 48 h. Strains and plasmid lists are provided in S5 Table.

### Growth curves

All growth curve experiments were performed in a Tecan SPARK 10M microplate reader using polystyrene flat-bottomed, non-treated, sterile 96-well plates. Overnight cultures grown in LBMC were washed and were subcultured (1:100 dilution) in MOPS-GS medium. The plates were programmed to continuously shake at 150 rpm. and the temperature was maintained at 30 °C. Optical density was measured at 600 nm every 60 min.

### Peptides

All chemically synthesized peptides were purchased from Genscript. The purity of all peptides was >99% and verified by high-performance liquid chromatography. Mixture of all regioisomers were used for all peptides as previously described [10].

### Antimicrobial Activity Assays

NCR247 sensitivity of *S. meliloti* and *E. coli* cells was determined as described previously [43] with modifications using early exponential phase cultures in 3-(N-morpholino) propanesulfonic acid (MOPS) buffered minimal medium (50 mM MOPS, 1 mM MgSO_4_, 0.25 mM CaCl_2_, 19 mM glutamic acid, 0.004 mM biotin, pH 7.0) Mops-GS supplemented with 1% casaminoacid.

### Scanning Electron Microscopy

SEM analysis was performed as described previously [30] with slight modifications. Untreated or 15 µM L- or D-NCR247 treated cells were fixed with 2% (vol/vol) glutaraldehyde and 3% paraformaldehyde in cacodylate buffer (0.05 M, pH 7.2) overnight. Cells were dehydrated serially in 50%, 75%, 90%, 95% and 3×100% ethanol. Then, the cells were dried in a critical point dryer, mounted on a stub, and sputter-coated with a 10 nm gold coating, and then viewed in a SEM (JOEL JSM-7100F/LV).

### ICP-MS

For bacterial samples, 1 ml of sample was spun down, and the pellet was resuspended in 40 μl of 100% HNO_3_ and heated at 98 °C for 1 h. The supernatant of the solution was mixed with metal-free water to make it up to 2 ml and ICP-MS analysis was performed as previously described [12]. The same number of cells were spun down for protein analysis using the BSA method and data were normalized to the amount of protein in each sample. Agilent ICP-MS instrumentation with MassHunter 4.4 was used to collect data.

### RNA isolation and RT–qPCR analysis

Cells were grown in LB medium until they reached an OD600 nm of 0.2. Then cells were spun down and suspended in MOPS-GS and synchronized according to previous method. Then cells were then treated with respective peptides and recovered at 30 mins for FeuP/FeuQ, ExoS/ChvL, RirA related genes and 90 mins for ctrA related genes. NCR247 was then added and 5 ml of appropriate cultures were spun down at given time intervals. Total RNA was extracted using the TRIzol (Thermo Fisher Scientific) method. A Qiagen RNeasy kit was used to purify the RNA. On-column DNA removal was carried out using DNase I from NEB. A total of 500 ng of each RNA sample was used to make complementary DNA using an iScript cDNA synthesis kit (Bio-Rad). RT–qPCR assays were performed as previously described [12]. The standard curve method was used for relative quantification. In brief, a standard curve was generated for each gene of interest (including sm*c00128*) by setting up qPCR reactions to amplify increasing amounts of *S. meliloti* Rm1021 genomic DNA. All the primer sets used resulted in a proportional dose– response curve with *R*2 > 0.99, which confirmed their efficiency. This curve was then used for extrapolating the relative expression level of each gene of interest in a particular sample to obtain the starting quantities (SQ). This value was then normalized to the SQ values of s*mc00128* obtained for the respective samples. These normalized values were then expressed as an average of triplicates, with s.d. represented by the error bars.

### *In vitro* translation assay

In vitro translation reactions were performed with the “Pure Express in Vitro Protein Synthesis” (Invitrogen Cat no: K990100) in the presence of different concentrations of NCR247 variants as described previously [11]. GFP fluorescence was measured in a Tecan Plate reader *(λ_ex_ = 485 nm, λ_em_ = 530 nm)* after 30 mins. In the vector used (a gift from the Baker laboratory, Massachusetts Institute of Technology), production of the GFP was controlled by the T7 promoter.

### Western Blot

Western blot was performed on the reactions from *in vitro* translation assay. 5 µl of the product was loaded. Anti-GFP antibody (ab6556) was used as primary at a dilution of 1:1000 and Goat anti -rabbit secondary (ab205718) was used at dilution of 1:10000.

### Biolayer Interferometry

Biolayer interferometry was carried out using ForteBio Octet RED96 biolayer interferometer following the manufacturer’s instructions for a standard kinetic assay. Biotinylated L- or D-NCR247 was loaded on to streptavidin coated biosensor tips. Peptides incubated in 200 μl assay buffer (water (pH 7.4)), each for 60 s. was loaded onto each biosensor tip at the defined concentration until the binding signal reached a value of >1.4. Biosensor tip loading was followed by incubation in assay buffer for 60 s. Association between the ligand and the purified SbmA analyte was observed over a time frame of ∼116 s in assay buffer. To stop binding kinetics for dissociation, the biosensor tips were placed back into an assay buffer not containing any analyte for 1200 s.

#### Pulldown assay

Log phase grown *S. meliloti* were lysed using French press homogenizer and split into three aliquots. 10 µM of N-terminal Biotin labelled L- or D-NCR247 was added to the extracts and incubated at room temperature for 3 hours. Pull down was performed using Dynabeads™ MyOne™ Streptavidin T1s (Invitrogen) on a magnetic rack. After washing 3 X with PBS. Equal amount of SDS loading buffer was added and boiled before loading to 12% SDS PAGE gel. Standard silver staining was then performed.

## Acknowledgments and funding sources

We thank the Walker lab members for helping with various aspects of preparation of this manuscript. We thank Dr. Joel Griffitts for the plasmid pJG206. This work was supported by the National Institutes of Health (NIH) Grant RO1 GM031010 (to G.C.W.) and funds from the Stowers Institute for Medical Research (to S.S). G.C.W. is an American Cancer Society Professor.

## Notes

### Competing Interest Statement

The authors have declared no competing interest.

